# miRNA Biomarkers in Prostate Cancer: Leveraging Machine Learning for Improved Diagnostic Accuracy

**DOI:** 10.1101/2024.10.25.618146

**Authors:** Shweta Singh, Abhay Kumar Pathak, Sukhad Kural, Lalit Kumar, Madan Gopal Bhardwaj, Mahima Yadav, Sameer Trivedi, Parimal Das, Manjari Gupta, Garima Jain

## Abstract

Prostate cancer (PCa) diagnosis often relies on prostate-specific antigen (PSA) testing, but its high false-positive rates often lead to unnecessary biopsies. MicroRNAs (miRNAs) have emerged as promising non-invasive biomarkers for cancer detection due to their stability in biological fluid and disease specificity. Despite their potential, the clinical translation of miRNAs as non-invasive cancer biomarkers is hindered by several challenges - population-based variability, environmental Factors, methodological Inconsistencies, lack of standardization, normalization Issues, and complexity of the biological System. These factors significantly impact the consistency of miRNA expression readouts, particularly in terms of Ct-values, across different studies, which in turn affects the determination of cutoff values that are crucial in a diagnostic setup. This preliminary study offers a pilot demonstration for integrating miRNA biomarker expression with machine learning (ML), which can help identify patterns and improve classification, potentially reducing the reliance on fixed cutoff values in certain contexts and pave the path to wider clinical translation.

We analyzed the expression of key miRNAs (miR-21-5p, miR-221-3p, and miR-141-3p) in blood samples from patients with PCa and benign prostatic hyperplasia (BPH). Utilizing a Random Forest classifier, we achieved an accuracy of 77.42%, a precision of 86.21%, a recall of 71.43%, and an AUC-ROC score of 0.78. The application of ML enabled us to leverage complex features, such as combinations and ratios of miRNA expression data, which enhanced the robustness and reliability of the diagnostic model. Additionally, bioinformatics analysis of the preferential features identified by the ML model confirmed the biological relevance of these miRNAs in PCa-related pathways, further supporting their potential as clinical biomarkers.

In the future, ML is poised to significantly enhance diagnostic performance compared to traditional linear analyses of a limited set of biomarkers. While our study did not explore multiple populations or the effects of methodological variables, it highlights the potential of ML by demonstrating improved accuracy and eliminating the need for cutoff values. This capability could broaden the applicability of miRNA-based diagnostics, making them more reliable and actionable in clinical settings. However, to fully realize this potential, further validation with larger and more diverse cohorts is essential. Overall, this study lays the groundwork for utilizing ML-enhanced miRNA panels as powerful tools for the early and non-invasive diagnosis of PCa in future clinical practice.

**Graphical Abstract:** 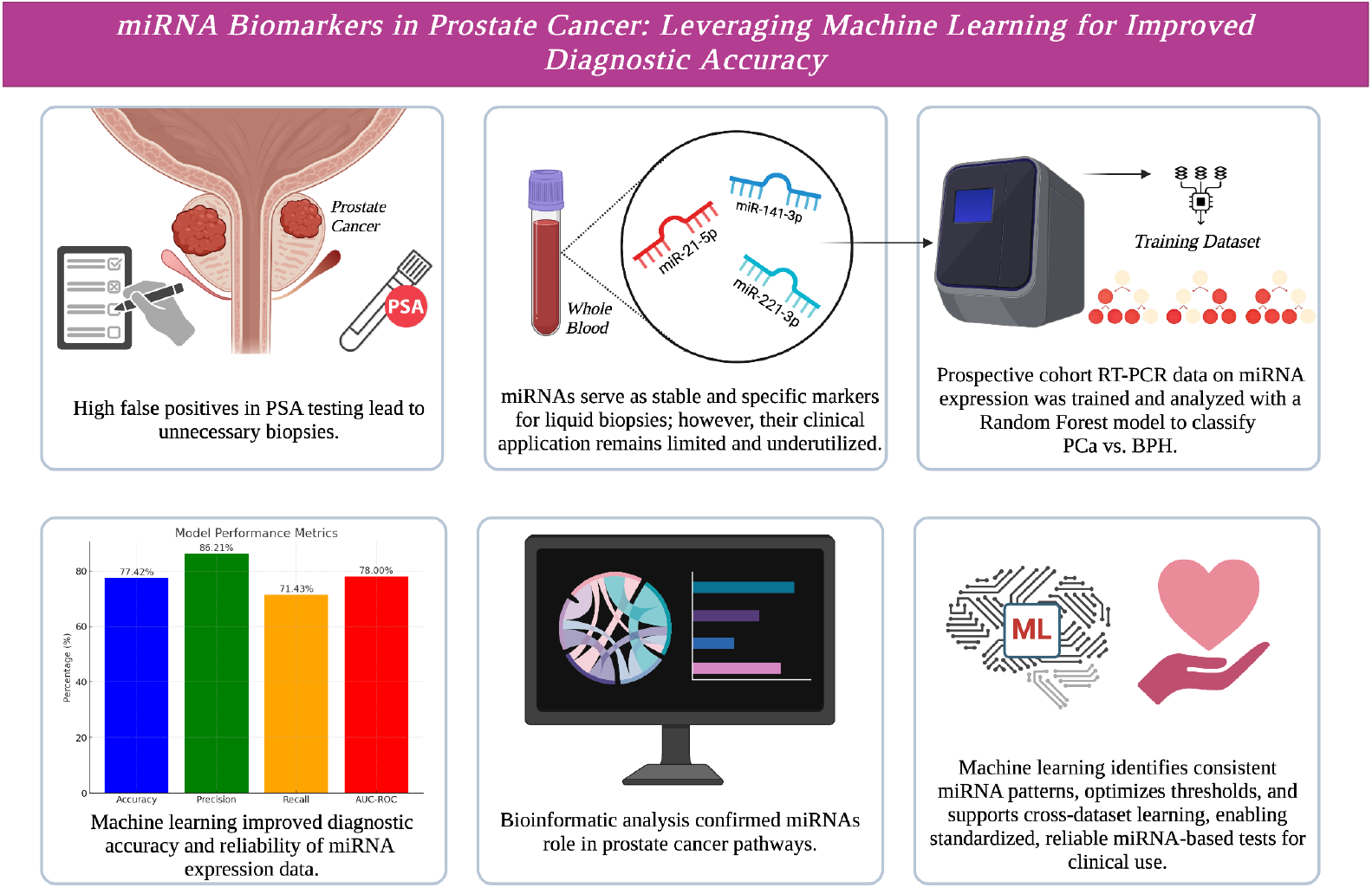

## 1. INTRODUCTION

MicroRNA (miRNA) biomarkers have emerged as critical players in disease diagnostics, particularly in cancer research over the past decade. Known for their stability and differential expression in disease versus healthy subjects, miRNA biomarkers are promising candidates for early cancer diagnosis and monitoring **(1,2)**. The advent of high-throughput technologies like Next-Generation Sequencing (NGS) has revolutionized miRNA biomarker discovery, providing comprehensive datasets that reveal miRNA expression profiles across various disease conditions **(3)**. Despite these advancements, the clinical application of miRNA biomarkers remains limited **(4,5)**, primarily due to the high cost and complexity of NGS, which makes it unsuitable for routine diagnostics in resource-limited settings **(6)**. Consequently, more affordable and accessible technologies such as Reverse Transcription Polymerase Chain Reaction (RT-PCR) has become the primary tools for validating discovered biomarkers **(7,8)**. However, the diagnostic application of miRNA biomarkers using RT-PCR-based methods faces significant challenges **(9)**. Current approaches for miRNA biomarker validation vary widely in terms of experimental design, internal controls, sample preparation, and data analysis **(10)**. A major concern is the variability introduced by differences in devices, methodologies, and sample handling. Such inconsistencies can result in data variations that do not accurately reflect the biological state of the patient but instead stem from technical artifacts. This lack of uniformity undermines the reliability of miRNA-based diagnostics in clinical settings. While PCR-based methods are sensitive for targeted miRNA quantification, they still exhibit relatively lower sensitivity compared to high-throughput approaches like NGS **(11)**. Variability and low sensitivity has hindered the translation of promising research findings into clinical trials and diagnostic applications. Furthermore, the necessity for large-scale and prospective clinical studies for validation remains indispensable, posing significant resource and ethical challenges. Addressing these challenges is essential for improving diagnostic accuracy and enhancing the clinical utility of miRNA biomarkers.

Machine Learning (ML) is a subset of artificial intelligence (AI) that enables systems to automatically learn and improve from experience without being explicitly programmed. ML models use algorithms to identify patterns in large datasets, making predictions or decisions based on that information. These models can be supervised (trained on labeled data), unsupervised (finding patterns in unlabeled data), or semi-supervised and reinforcement learning (which involve learning from feedback). Over time, the model’s performance improves as it processes more data, making ML a powerful tool in various fields, including healthcare **(12)**.

Linear methods, such as linear regression, assume a direct relationship between inputs and outputs. These models can struggle when data shows high variance or complex relationships, leading to underfitting or overfitting. ML models, especially those like decision trees, random forests, and neural networks, are more flexible and equipped to handle variance **(13)**. They capture non-linear relationships and interactions between features more effectively than linear models. For example, ensemble techniques such as Random Forests use multiple decision trees to reduce variance by averaging predictions, making ML models robust in noisy and complex datasets where traditional linear models would fail **(14)**.

ML models are revolutionizing biological research by analyzing complex datasets like genomics, proteomics, and medical imaging. In biology, datasets often exhibit high dimensionality, non-linearity, and noise, where ML excels **(15)**. For instance, ML models have extensively been used in biomarker discovery, such as in cancer diagnostics, by analyzing RNA sequencing or microarray data **(16)**. Supervised learning models, which excel in classifying labeled data, however remain underutilized in the context of RT-PCR data. To the best of our knowledge, very few studies have utilized RT-PCR data to effectively leverage supervised learning for enhancing the diagnostic accuracy **(17,18)**. By training ML models on a large set of patient-specific RT-PCR data, ML models can capture subtle patterns in multiple miRNA expressions that may go undetected by traditional analysis methods, to improve the sensitivity (the ability to accurately identify true positives) and specificity (the ability to accurately identify true negatives) of miRNA biomarkers.

The aim of our study was to assess the expression levels of a miRNA panel in the blood of Prostate Cancer (PCa) patients compared to those with benign prostatic hyperplasia (BPH) and apply supervised ML algorithms to enhance the sensitivity and specificity of these biomarkers. By leveraging the strengths of ML alongside the cost-effectiveness and accessibility of RT-PCR, we aimed to not only improve biomarker panel performance but also bridge the gap between discovery research and clinical application.

PCa is one of the most prevalent cancers in men over the age of 60. The standard screening method is the serum prostate-specific antigen (PSA) test. However, while PSA levels often rise in prostate-related conditions, it is not specific to PCa **(19)**, leading to high false-positive rates, overdiagnosis, and unnecessary biopsies. The gold standard for distinguishing PCa from BPH remains a transrectal prostate biopsy, which is invasive and uncomfortable, causing many patients to avoid it **(20)**. Since PCa and BPH differ significantly in their life-threatening risk, treatment, and intervention strategies **(21)**, early differentiation is crucial for informed decision-making. Therefore, there is an urgent need for more accurate, non-invasive diagnostic tools to distinguish between these conditions.

The discovery of miRNAs in the plasma of PCa patients sparked significant interest in their potential as liquid biopsy biomarkers for PCa **(22)**. Numerous studies have identified and validated various miRNAs as potential biomarkers; however, the results have been somewhat inconsistent. Different studies have reported varying miRNA panels, with conflicting data on expression levels, leading to mixed conclusions about their effectiveness **(23,24)**.

Based on previous studies, we selected a group of promising miRNAs for investigation in our study, as outlined in the methods section: miR-21-5p, miR-141-3p, miR-221-3p, miR-93-5p, miR-19b-3p, and miR-375-3p. While urine, plasma, and serum are commonly used samples in most of the studies **(25,26)**, whole blood also presents a good source of biomarkers with distinct advantages, including a higher miRNA yield, reduced susceptibility to technical errors, and a more comprehensive representation of the systemic profile **(27)**. In this study, we focused on miRNA profiling from whole blood using the RT-PCR method, conducted in three distinct phases with separate cohorts. In the discovery phase, the most significant miRNAs were identified from the discovery cohort. The verification phase involved using RT-PCR data from the verification cohort to train the ML model. In the final validation phase, the ML model was tested on a completely new dataset. **Fig. 1** shows all the stages of the study schematically. The results showed that combining ML with miRNA biomarkers significantly improved diagnostic efficiency compared to using individual miRNAs alone. This integration of ML and RT-PCR offers a more accurate, cost-effective, and accessible diagnostic tool, with the potential to transform early cancer diagnosis and management in clinical practice.

**Fig. 1:**
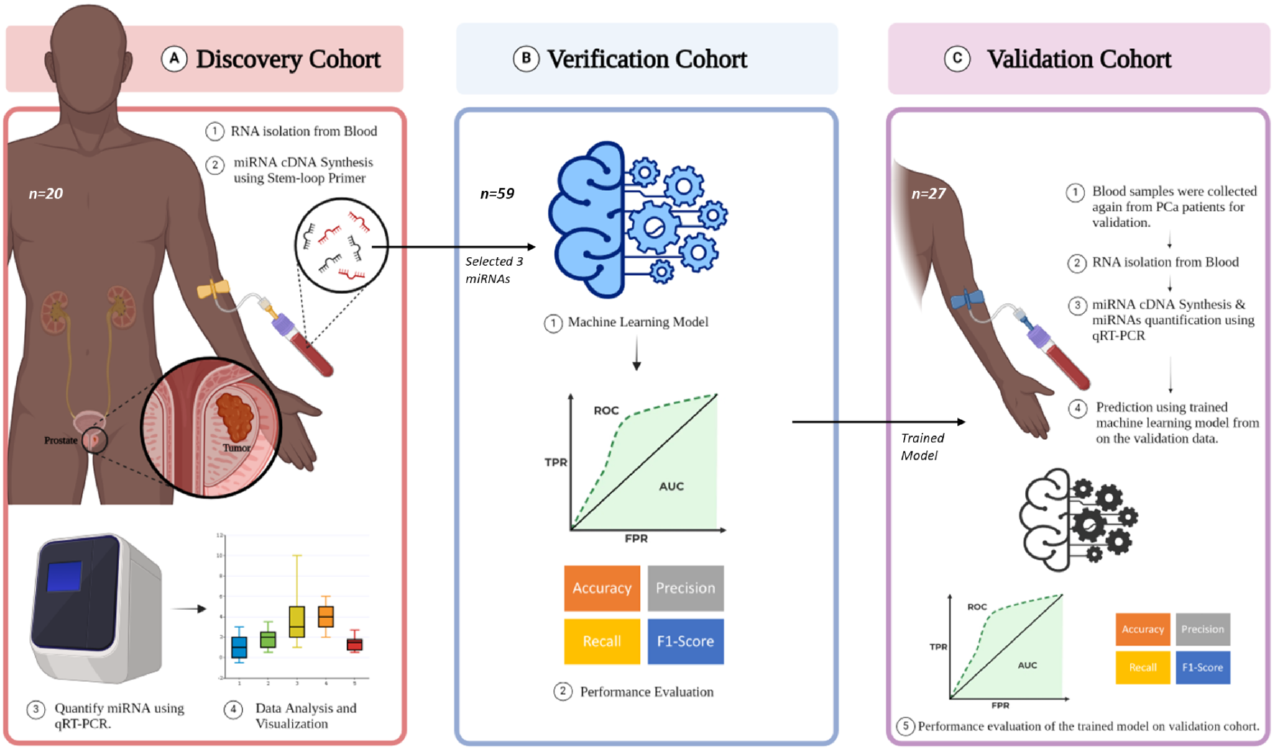
Workflow of the Study Design **(A)** Discovery Cohort; **(B)** Verification Cohort; **(C)** Validation Cohort. Figure created using Biorender (https://biorender.com/)

## 2. METHODS AND MATERIAL

### 2.1. Study population

We conducted a prospective cohort study of 86 participants from November 2022 to August 2024 in collaboration with the Centre for Genetic Disorders and the Department of Urology at the Institute of Medical Sciences, Banaras Hindu University. The study was approved by the ethical committee of the Institute of Science at Banaras Hindu University (ECR/226/Indt). The research investigated a treatment-naive cohort of 81 patients, clinically suspected of having PCa were planned for a prostate biopsy, based on digital rectal examination (DRE) and elevated PSA levels illustrated in **Table 1**; along with 5 healthy subjects. Exclusion criteria involved: (a) lack of complete medical records; (b) non-urothelial malignancies; (c) other urological diseases (e.g., urinary tract infections); (d) refusal to consent; and (e) prior androgen deprivation therapy (ADT) or radiation therapy. In the second phase, de-identification of collected samples was performed, followed by classification into two groups based on histopathological examination of 12-core prostate biopsies resulted in group 1 PCa (n=51) and group 0 combination of BPH (n=30), healthy (n=5).

**Table 1:**
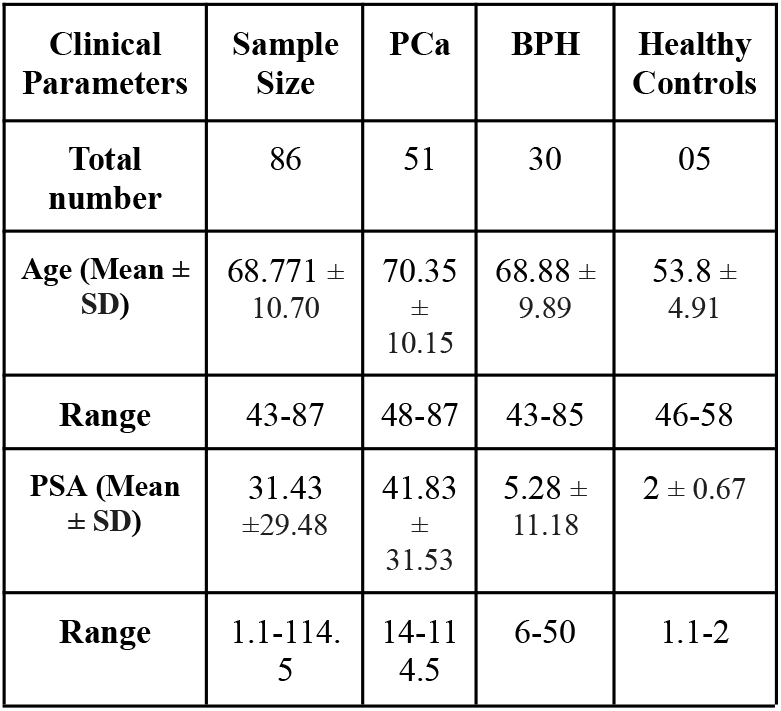
Clinicopathological characteristics of prostate biopsy patients.

The group 1 (51 PCa samples) could be categorized into three distinct risk groups. On the basis of Gleason score as presented in **Table 2**. This distribution underscores a predominance of intermediate and high-grade cases within the cohort studied.

**Table 2:** Stratification of PCa sample by Cancer Grade, Gleason Score, and Risk Categories.

| Cancer Grade | Grade Group | Gleason Score | No. of PCa samples |
| --- | --- | --- | --- |
| Low to very low | Grade Group 1 | 3+3 | 4 |
| Intermediate | Grade Group 2,3 | 7(3+4) , 7(4+3) | 20 |
| High to very high | Grade Group 4,5 | 8, 9-10 | 27 |

### 2.2. MicroRNA panel selection

A systematic literature review was conducted in PubMed to find microRNAs implicated in PCa initiation, progression and metastasis using the terms ‘prostate cancer’ and ‘microRNA’. The cited references of studies were checked for further articles of interest. All articles including original articles, reviews and abstracts were considered. Based on our review we chose a panel of 6 miRNAs for profiling in whole blood, all of which had at least 2 independent studies published showing diagnostic capacity of miRNA with preference given to those with concomitant supporting mechanistic data mentioned in **Supplementary Table 1**.

### 2.3. Blood collection, storage and RNA isolation

Peripheral venous blood was collected from the patient into collection tubes coated with EDTA prior to the prostate biopsy. Total RNA was extracted from 400 µL of blood using 750 µL of Trizol reagent (Cat No. 15596026) subjected to disruption and homogenization. Following this,200 µL of chloroform was added, the mixture was vigorously shaken, and then incubated at 4°C. After centrifugation at 12,500 × g for 15 minutes at 4°C, the sample was partitioned into three distinct phases. The upper aqueous phase was carefully separated, combined with 500 µL of isopropanol, and centrifuged at 10,000 × g for 10 minutes at room temperature. The supernatant was removed, the pellet was air-dried, and resuspended in 20 μL of DEPC-treated water before being stored at -80.The concentration and quality of the extracted RNA were assessed using NanoDrop (Thermo Scientific, USA).

### 2.4. Reverse Transcription & RT-PCR

The RevertAid First Strand cDNA Synthesis Kit (Catalog #K1622, Thermo Scientific, USA) was used for the reverse transcription of miRNA samples. This process involved a reaction mixture comprising a miRNA-targeted stem-loop primer for reverse transcription. The details of primer sequence have been given in **Supplementary Table 2**. For quantitative PCR (RT-PCR), reactions were carried out in the Applied Biosystem QuantStudio 6 Flex Real-Time PCR System (Thermo Scientific, USA). The reaction mixture comprised 1μL of pre-amplified cDNA (1:5 pre-diluted), 6.25μL of Maxima SYBR Green/ROX RT-PCR Master mix (2X, Catalog #K0221, Thermo Scientific, USA), miRNA-specific 1μL forward primer and 1μL reverse primer 10μM, and nuclease-free water to make up the final volume of 12.5μL. The amplification program involved an initial temperature of 95°C for 10 min, followed by 40 cycles of 95°C for 15 s and 60°C for 1 min. Ct values were obtained by finding where the threshold line intersects with the amplification curve. Delta Ct values were then calculated by subtracting the Ct value of the endogenous control, RNU6, from the Ct value of the target gene for each sample.

### 2.5. Study Design

The study was designed in three cohorts: discovery, verification, and validation presented in **Fig. 1**. The discovery cohort comprised 20 samples, including BPH (n=9) and PCa (n=11) cases, and was used to analyze the expression patterns of seven selected miRNAs using RT-PCR. The hypothesis test (Mann-Whitney U) was employed to identify the most differentially expressed miRNAs among the selected group, resulting in the selection of three miRNAs: miR-21-5p, miR-141-3p, and miR-221-3p. In the verification cohort (n=59) composed of BPH (n=27) and PCa (n=32), the expression data for these 3 selected miRNAs was collected and this data was formed to build and train a ML model. Finally, the validation cohort (n=27): BPH (n=11) PCa (n=16) with the sample that were previously unexposed to the trained model. The aim of the validation cohort was to evaluate its performance of the trained ML model.

### 2.6. ML-Model Development

#### 2.6.1. Preprocessing

For the preprocessing step, NULL values have been imputed by the median value. All the operations have been performed on Delta Ct values.

#### 2.6.2. Statistical analysis

For the hypothesis test we have used Mann-Whitney U test, also known as the Wilcoxon-sum test, is a nonparametric test that is used to compare the differences between two independent classes, because its non-assumption of normal distribution is a versatile tool to use. ROC(Receiver Operating Characteristic) curve is a graphical representation that shows the performance of a binary model at different threshold settings. Using two values of TPR (True Positive Rate) and FPR (False Positive Rate) it plots two parameters at different threshold levels. The AUC area under the ROC curve quantifies the overall ability of the model to differentiate between negative (BPH) and positive (PCa) classes. Hypothesis test and ROC-AUC used for preliminary selection of differentially expressed biomarkers from the panel.

#### 2.6.3. Feature Selection

Initially the study started with seven biomarkers for the classification of benign and malignant PCa aiming to show differentially expressed patterns in both categories. Each biomarker is evaluated based on Mann-Whitney U test to identify most significant biomarkers in terms of discriminative power in both classes.

#### 2.6.4. Machine Learning

In this study, Random Forest has been employed as a powerful classifier, combining an ensemble of decision trees to capture the complex patterns of RT-PCR data. The detailed steps used in the modeling shown in **Fig. 2**. Each tree within the ensemble is constructed from bootstrapped samples and guided by a randomly selected subset of features, which helps to reduce the correlation between trees and enhances the overall diversity of the model. By working collectively, these decision trees independently vote on each diagnosis, minimizing the influence of any single outlier and improving the robustness of the final classification. Through this approach, Random Forest efficiently handles non-linear relationships within the dataset, as each tree contributes to the model’s capability to generalize. The randomness introduced in both sampling and feature selection mitigates overfitting, leading to improved stability and accuracy.

**Fig. 2:**
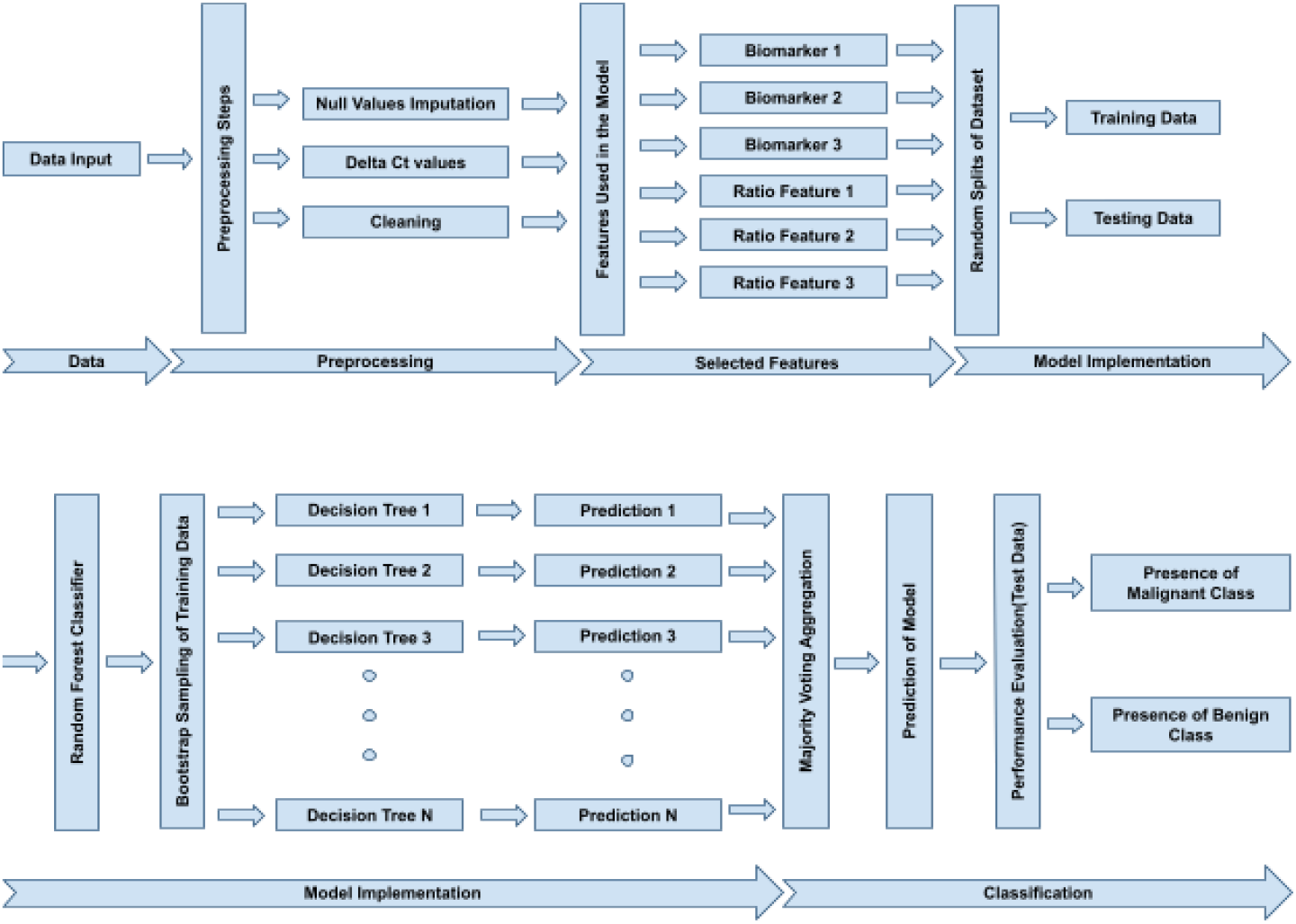
Workflow of the proposed ML model in the verification cohort.

Ultimately, the aggregate voting of these trees forms the final prediction, leveraging the “wisdom of the crowd” to enhance decision-making in PCa classification.

### 2.7. Bioinformatics Analysis

To predict the target genes of the 3 miRNAs, we used the Database like miRTarBase and miRBase in September, 2024 **(28,29)**. Target gene lists from both the sources were combined by removing any duplicates. All selected target genes were listed in a common list referred to as miRNA Target Gene Panel (mTGP). This list was used to analyze functional aspects by Gene ontology based functional analysis using ShinyGO tool **(30)**. A PCa specific gPCRene panel was created using GeneCards, KEGG and String database **(31)**. We employed the keyword “PCA” (Prostate Cancer) to retrieve gene entries from the databases to ensure the inclusion of genes associated specifically with PCa. All the genes reflected from the search were included in the PCa Gene Panel (PGP). To improve the reliability of target gene results, we identified the overlapping target genes from mTGP and PGP by using the VennDiagram package of R software. Then, these overlapping target genes were used to study protein-protein interaction and cancer context specific network construction using PINAv3.0.

## 3. RESULTS

### 3.1. miRNA Expression Analysis in Discovery Cohort

In the discovery cohort, miRNA expression analysis was done using RT-PCR, in both BPH and PCa groups. The Mann-Whitney U test showed that all 6 miRNAs had p-values below 0.05, indicating significance of selected miRNAs. To ensure the use of most significant feature in verification phase, miR-21-5p, miR-221-3p, and miR-141-3p were selected exhibiting the most significant differences, with p-values of 0.000224, 0.000153, and 0.004293, respectively. Additionally, miR-375-3p, miR-19b-3p, and miR-93-5p also displayed significance with p-values ranging from 0.004747 to 0.037008 **(Fig. 3, Table 3)**. These findings, illustrated by the boxplots of ΔCt values, highlight the potential of these miRNAs as diagnostic biomarkers for PCa. Based on these results, the three most significant miRNAs were selected for further study in the verification cohort.

**Table 3:** P-values of different genes with respect to their classes of BPH and PCa using Mann-Whitney U test.

| Gene Number | Gene Name | P-Value |
| --- | --- | --- |
| 1 | miR-21-5p | 0.000224 |
| 3 | miR-375-3p | 0.004747 |
| 4 | miR19B-3p | 0.017574 |
| 5 | miR-93-5p | 0.037008 |
| 6 | miR-141-3p | 0.004293 |
| 7 | miR-221-3p | 0.000153 |

**Fig. 3:**
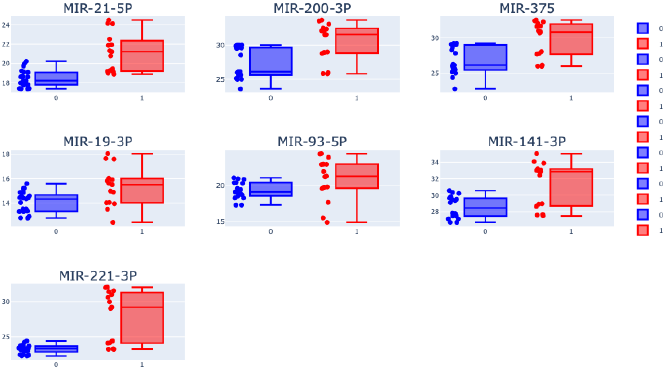
Boxplot showing the Delta Ct value expression in different miRNA’s. Where ‘0’ represents BPH ‘1’ represents the PCa.

### 3.2. ML Model Development on Verification Cohort

Similar to the discovery cohort, the verification cohort was used to generate expression data of selected miRNAs using RT-PCR. In the verification cohort, 6 features were created - 3 the individual miRNAs as features (miR-21-5p, miR-141-3p, miR-221-3p) and their ratio-based combinations (21_141, 141_221, and 221_21). The performance of each feature was assessed using ROC curve analysis demonstrated in **Fig. 4**. The resulting AUC scores ranged from 0.48 to 0.52, indicating that the both individual features and ratios performed poorly when used as independent features. This suggests that these miRNAs, when considered independently, do not possess strong discriminative power for distinguishing between BPH and PCa, limiting their standalone utility in diagnostic predictions.

**Fig. 4:**
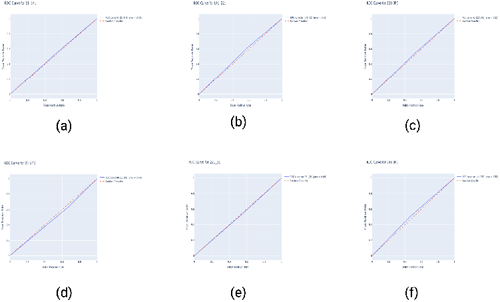
ROC graph with respected AUC score of selected biomarkers features for the Model **(a)**21_141 = dCt of miR-21-5p/ dCt of miR-141-3p; **(b)**141_221 = dCt of miR-141-3p/ dCt of miR-221-3p; **(c)** miR-221-3p; **(d)** miR-21-5p; **(e)** 221_21 = dCt of miR-221-3p/ dCt of miR-21-5p; **(f)** miR-141-3p.

In the verification cohort, a random forest model was applied using each miRNA expression data along with ratios in 3 combinations as features. The trained ML model showed an accuracy of 77.42%, precision of 86.21%, recall of 71.43%, and an F1 score of 78.12%, as illustrated by the confusion matrix presented in **Fig. 5 (a)**. The ROC curve for the random forest model showed an AUC of 0.78, reflecting a good balance between true positive and false positive rates shown in **Fig. 5 (b)**. This indicates that the combination of miRNA as features and supervised training ML-model can enhance the overall ability of the panel of biomarkers.

**Fig. 5:**
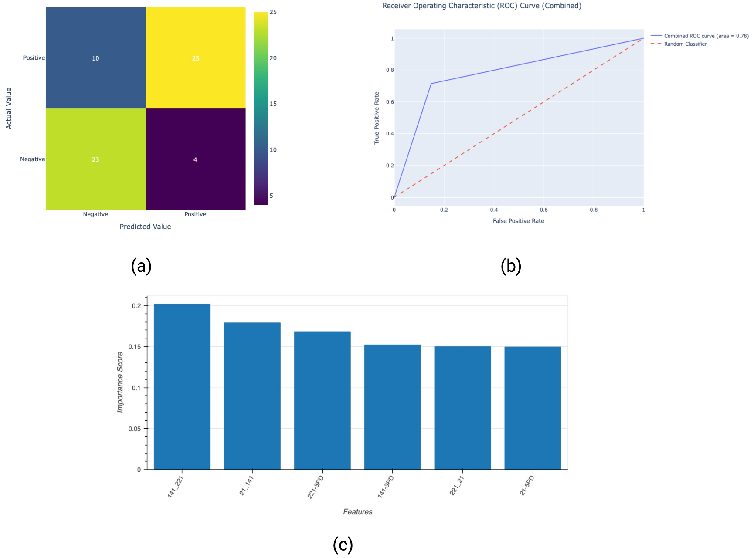
**(a)**Confusion matrix for the model trained with differentially expressed miRNAs on test set in verification cohort; **(b)** ROC AUC Graph; **(c)** Feature importance employed by Random Forest Model. 141_221 refers to expression ratio of miR-141-3p/miR-221-3p, 21_141 refers to expression ratio of miR-21-5p/miR-141-3p, and 221_21 refers to expression ratio of miR-221-3p/miR-21-5p.

The ensemble model, being composed of multiple decision trees, assigns importance to features based on their ability to improve classification accuracy. Further analysis of feature importance, generated by the random forest classifier, revealed miRNAs and feature combinations that contributed most to the model’s predictive accuracy. As shown in **Fig. 5 (c)** the feature importance plot, the ratio feature of miR-141-3p and miR-221-3P was the most significant contributor, followed closely by ratio of miR-21-3p and miR-141-3p. This indicates that these features have the strongest influence on the model’s predictions and are the key drivers behind its discriminative performance.

### 3.3. Validation of designed ML-model on Validation cohort

To validate the performance of the trained model, we used an independent cohort that was previously unexposed to the ML-model. When expression data from this validation cohort is employed to the trained ML-model it demonstrates significant performance across multiple evaluation metrics. The testing of the model showed an accuracy of 74.07%. It exhibited a precision of 71.43%, meaning it effectively identified true positive cases, while maintaining a recall (sensitivity) of 76.92%, which reflects the model’s capacity to capture most of the true positive instances of PCa. Furthermore, the model achieved a specificity of 74.07%, demonstrating its capability to correctly classify negative cases, effectively minimizing false positives shown in **Fig. 6 (a)**.

**Fig. 6:**
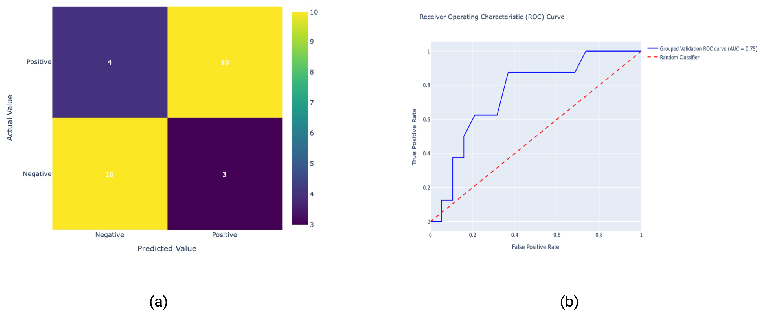
**(a)**Confusion matrix for the model trained with differentially expressed miRNAs on test set in validation cohort; **(b)** ROC AUC Graph for validation cohort.

The model’s ROC-AUC score of 0.75 confirms its discriminatory power, which is far better than current PSA based matrices **Fig. 6 (b)**.

### 3.4. Association of miRNA and their target genes with PCa pathogenesis

We predicted target genes for the selected miRNAs (miR-21-5p, miR-141-3p, and miR-221-3p), identifying 278 genes (mTGP panel). Using ShinyGO, we conducted Gene Ontology (GO) enrichment analysis to explore key biological processes (BP), cellular components (CC), and molecular functions (MF), presenting the top 20 in each category. The BP analysis revealed that the genes are primarily involved in regulating metabolic processes related to nitrogen compounds, macromolecules, and nucleobase compounds. In the MF category, the genes were linked to chromatin binding, DNA binding, and kinase activity. The CC analysis showed strong associations with ribonucleoprotein complexes, including the RNA-induced silencing complex (RISC) and transcription regulator complexes. Additionally, KEGG pathway analysis highlighted enrichment in cancer-related pathways, such as miRNAs in cancer, TNF, JAK-STAT, MAPK, and Akt signaling, as well as growth hormone and prolactin synthesis—processes relevant to cancer pathogenesis, as shown in **Fig. 7**.

**Fig. 7:**
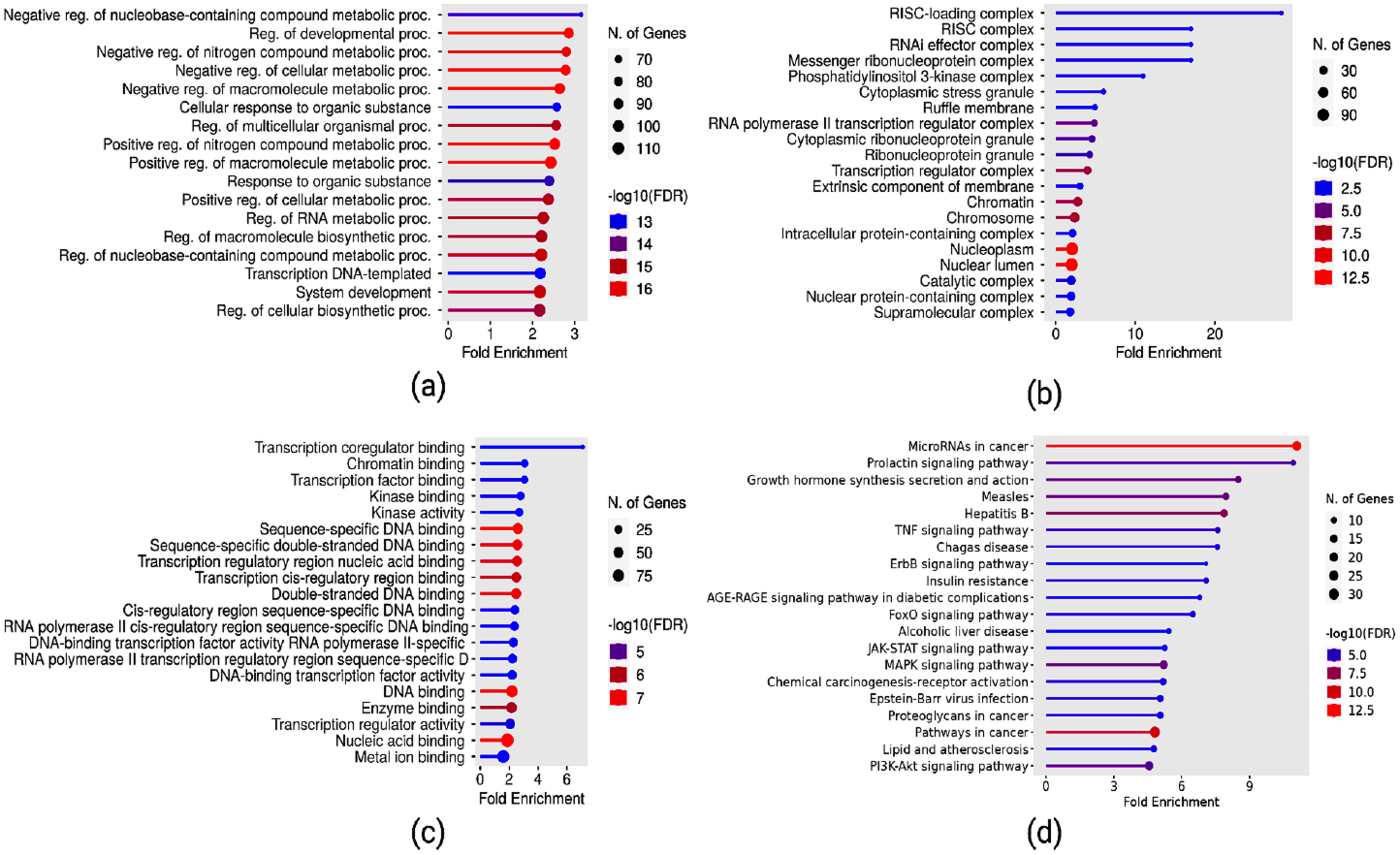
Gene Ontology and KEGG enrichment analysis of target genes for miR-21-5p, miR-221-3p,miR-141-3p. (a)Biological Processes:shows regulation of metabolic pathways crucial for cellular function and cancer progression (b)Cellular Component: reveals key roles in transcription factor binding and kinase activity. (c)Molecular Function : reveals key roles in transcription factor binding and kinase activity. (d)KEGG Pathway analysis highlights enrichment in cancer-related signaling pathways.

To determine which of the 3 miRNA could prominently be associated with PCa progression, identified target genes (mTGP) were screened against PCa gene panel (PGP), revealing 32 shared genes. Result of the chord diagram shows that all 3 miRNA interacted with at least one gene from PGP **(Fig. 8)**. The analysis highlighted miR-21-5p targeting the most PCa-associated genes, followed by miR-141-3p and miR-221-3p, suggesting these miRNAs are critical biomarkers for PCa progression.

**Fig. 8:**
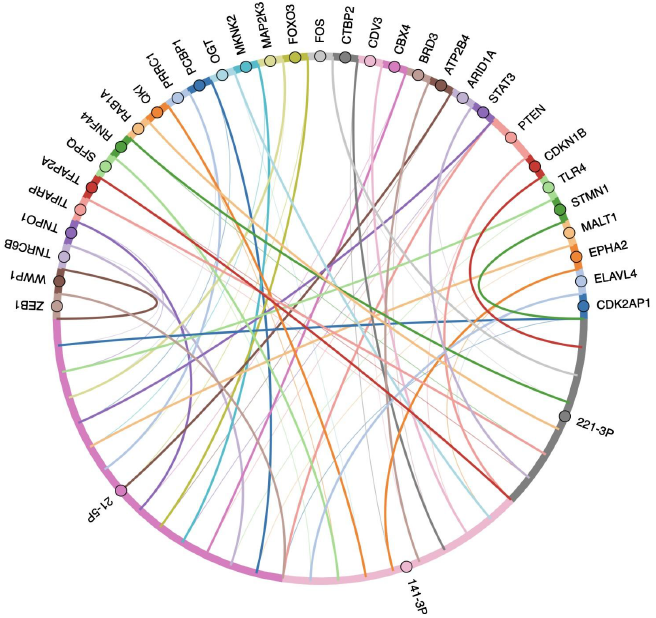
Chord diagram analysis of miR-21-5p, miR-141-3p, and miR-221-3p and their target genes associated with PCa.

KEGG pathway analysis on 15 genes out of 32 genes that interacted with miR-21-5p showed involvement of genes in general cancer related pathways such as PD-L1 and PD-1 checkpoint, Prolactin, HIF-1, NF-kB etc. Whereas combined analysis of 18 genes interacting with miR-141-3p and miR-221-3p showed more PCa specific pathways involvement such as Endocrine Resistance (**Fig. 9**).

**Fig. 9:**
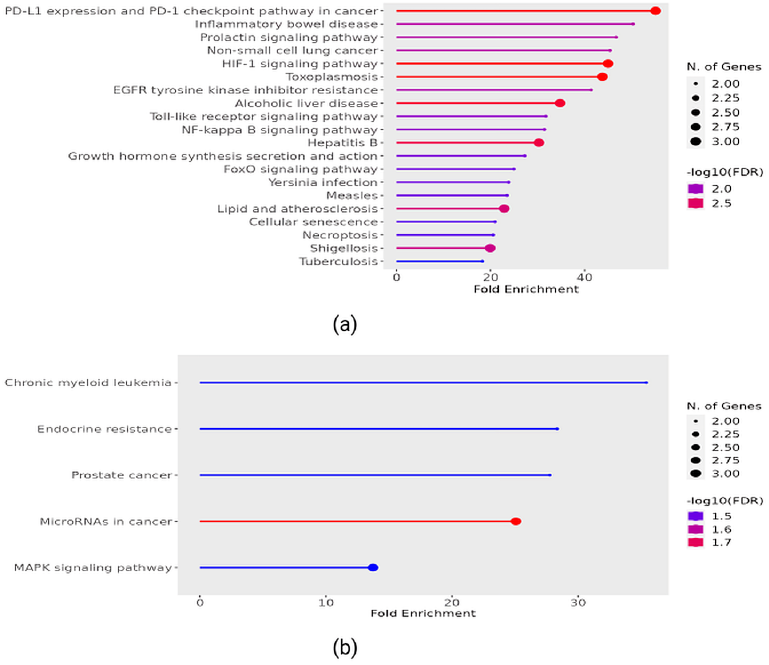
Pathway Enrichment Analysis of miRNAs and Their Target Genes in PCa. Top enriched pathways for target genes of **(a)** miR21-5p and, **(b)** for target genes of miR-141-3p and miR-221-3p.

Interestingly, the 32 identified genes are not independent but interact within cancer-specific protein-protein interaction (PPI) networks, as constructed using PINAv3.0 **(Fig. 10)**. This analysis revealed seven hub genes—RASA1 (RAS P21 Protein Activator 1), RHOB (Ras Homolog Family Member B), CDKN1B (Cyclin Dependent Kinase Inhibitor 1B), EPHA2 (EPH Receptor A2), ARID1A (AT-Rich Interaction Domain 1A), OGT (O-GlcNAc transferase), and CBX4 (Chromobox 4)—all associated with tumorigenesis. Specifically, RASA1 and ARID1A were strongly linked to the androgen receptor (AR) pathway, which is frequently disrupted in PCa. Other notable interactions showed connections between key oncogenes, such as SMAD2 (SMAD Family Member 2), TNFAIP1(TNF alpha induced protein 1), and CBX8 (Chromobox homolog 8), with these hub genes **(Fig. 10)**. A broader PPI network generated using STRING for the complete mTGP gene list identified additional hub genes like STAT3, PTEN, CDKN1B (Cyclin Dependent Kinase Inhibitor 1B), and FOS (), indicating these proteins play crucial roles in systemic regulatory networks **(Supplementary Fig. 1)**. Overall, the miRNA targets exhibited extensive interactions with commonly affected genes in PCa, underscoring their significant role in the development and progression of the disease through target gene regulation.

**Fig. 10:**
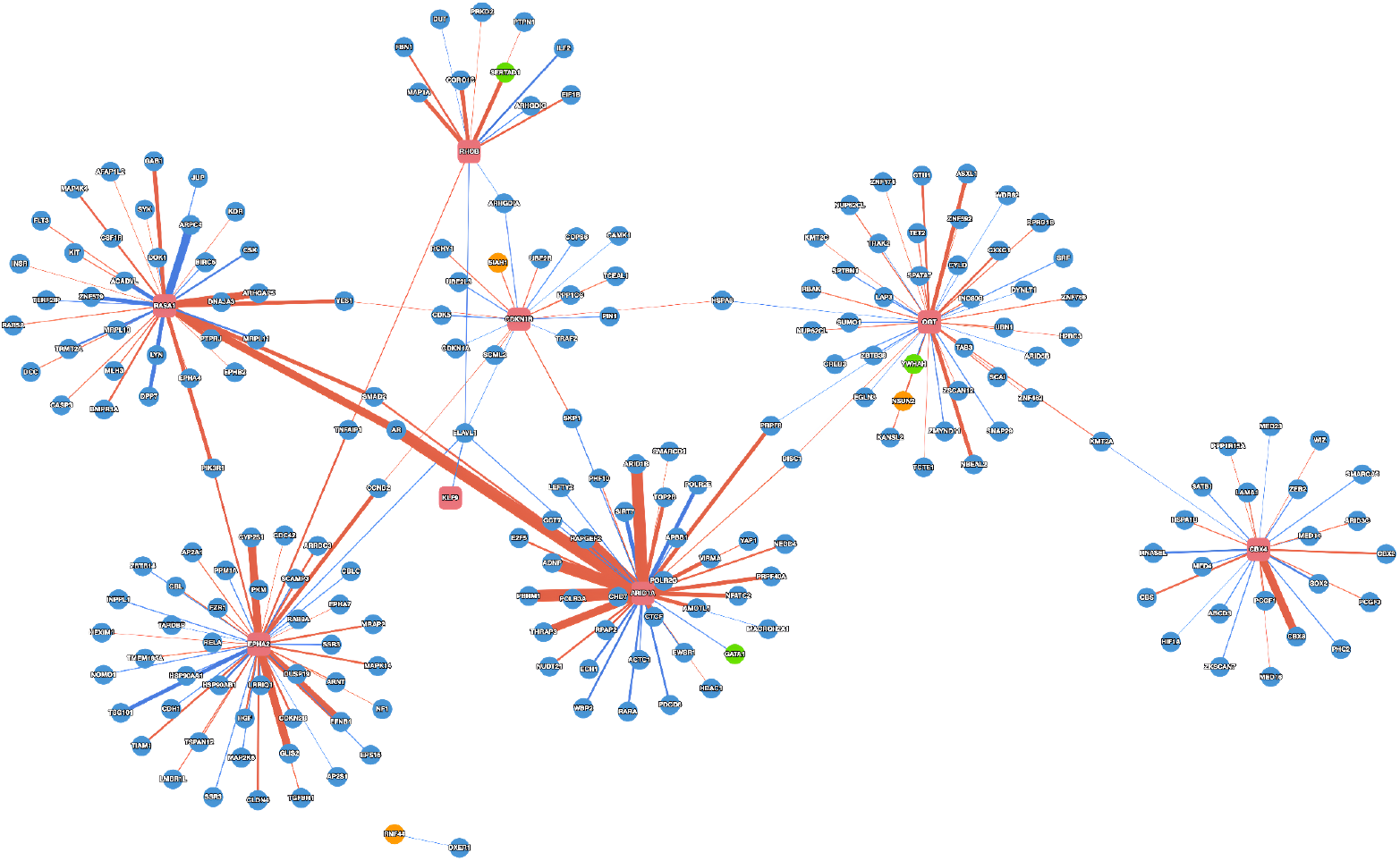
Protein-Protein Interaction (PPI) Network Highlighting Key Hub Genes in PCa. The key hub proteins are represented by red circles, exhibiting extensive interaction with proteins of both favorable (green circles) and poor (orange circles) prognosis related genes. Strong interactions demonstrated with thicker edges.

## 4. DISCUSSION

PCa diagnosis is often challenging, particularly in distinguishing it from BPH, which shares overlapping clinical features. The current reliance on PSA tests results in a high number of false positives, leading to unnecessary biopsies and patient anxiety leading to critical need for more reliable non-invasive biomarkers that can accurately differentiate between PCa and BPH **(32,33)**. miRNAs play a critical role in the progression and pathogenesis of PCa, and their use as biomarkers in various studies has yielded encouraging results **(34,35)**.

miRNAs circulate freely in the blood and reflect changes in tumor biology, making them ideal candidates for non-invasive diagnostic tools. However, based on inconsistencies in miRNA expression among various studies may warrant larger validation studies on diverse cohorts. Our study focused on analyzing expression of miRNAs with differential expression between PCa and BPH patients. We have selected 6 miRNA-miR-21-5p, miR-141-3p, miR-93-5p, miR-19b-3p, miR-375-3p and miR-221-3p - based on previous studies. Often referred to as an “oncomiR,” miR-21-5p is one of the most frequently upregulated miRNAs in various cancers, including PCa. It is known to promote tumor growth and progression by targeting tumor suppressor genes like PTEN, thereby contributing to cell proliferation, migration, and resistance to apoptosis **(36)**. miR-141-3p is a member of the miR-200 family and has been consistently shown to be upregulated in PCa patients, especially in advanced stages of the disease. It plays a role in epithelial-to-mesenchymal transition (EMT), a process crucial for cancer metastasis, and is also associated with androgen receptor signaling **(37)**. miR-221-3p is known for its role in regulating cell cycle and apoptosis, and is frequently overexpressed in PCa. It has been implicated in resistance to cancer therapies and is involved in modulating the expression of genes like CDKN1B (p27), which controls cell cycle progression **(38)**. miR-93-5p is associated with the regulation of tumor angiogenesis and has been found to be overexpressed in PCa **(39)**. miR-19b-3p is involved in oncogenic processes, including the suppression of tumor suppressor PTEN **(40)**. Unlike the others, miR-375-3p is often downregulated in PCa and acts as a tumor suppressor.

It has a role in regulating insulin secretion and has also been shown to influence cancer cell proliferation and migration by targeting genes like YAP1 **(41,42)**.

In the present study, the importance of the ML model as a tool to handle the subtle variation and identify complex patterns between different miRNA expressions was examined using PCa as a target disease and RT-PCR as a dataset. Random Forest classifier, the ML-model used in our study, is capable of identifying complex patterns in large datasets that may be overlooked by traditional statistical methods. By training our model on miRNA expression data, we significantly improved the diagnostic accuracy, achieving an AUC-ROC score of 0.78, which is a marked improvement over the single miRNA as biomarker and traditional PSA-based testing.

We chose to use whole blood as our biological sample for miRNA profiling, due to multiple reasons like higher yield of miRNAs compared to plasma or serum, comprehensive reflection of the systemic pathologies and capturing miRNAs from various tissues, including distant inflammatory sites **(43)**. However, a potential limitation is the complexity of whole blood samples, which contain miRNAs from multiple cellular sources, making it more challenging to attribute changes in miRNA levels specifically to a certain cancer progression. Interestingly promising results in cancers, like breast, pancreatic, and lung cancers have been reported using the use of miRNA profiling in whole blood yet for PCa this approach has received limited attention **(44–46)**. While other studies utilizing plasma and serum miRNAs have also reported high diagnostic accuracy, the results across studies tend to vary **(47,48)**. This is largely due to differences in study populations, sample processing techniques, and the lack of a standardized cutoff for Ct values in RT-PCR analysis. Instead of relying on fixed thresholds, ML models can adaptively determine optimal cutoff values based on data distribution, allowing them to tailor sensitivity and specificity to the specific cohort being analyzed. The ML model dynamically adjusts to these variations, offering a more flexible and robust approach to diagnosis, which traditional methods may not provide.

One challenge in the clinical use of ML models is their “black box” nature, which can make it difficult for clinicians to interpret results. To address this, we integrated bioinformatics analyses to confirm the biological relevance of the key features selected by our ML model. For instance, the importance of specific miRNA pair ratios miR-141-3p/miR-221-3p and miR-21-5p/miR-141-3p identified by the model was validated through their known roles in PCa progression, supported by target gene analysis. A chord diagram revealed that miR-21-5p had the most interactions with PCa-related genes, reinforcing its critical role in cancer development. This aligns with existing studies linking miR-21 to tumor growth, metastasis, and resistance to therapy. Additionally, KEGG pathway analysis showed that miR-21-targeted genes are involved in several cancer-related pathways, including the PD-L1/PD-1 checkpoint, prolactin, HIF-1, and NF-κB signaling pathways, which are known to regulate immune evasion, angiogenesis, and inflammation in cancer **(49,50)**. In contrast, the combined analysis of genes targeted by miR-141-3p and miR-221-3p highlighted their involvement in more PCa-specific pathways, particularly those related to endocrine function. These pathways play a key role in the hormonal regulation of PCa, such as androgen signaling. This finding suggests that miR-141-3p and miR-221-3p may have distinct roles in the progression of PCa, especially in hormone-sensitive and castration-resistant prostate cancer (CRPC). This explainability not only strengthens the trust in our model but also sheds light on the underlying biology of PCa.

We evaluated the performance of the trained ML model using a completely new dataset, which had not been previously used during training or testing. This step was crucial to prevent any potential data leakage, ensuring that the model’s performance reflects its ability to generalize rather than simply memorizing patterns from the training data. By testing the model on this fresh dataset, we aimed to assess its real-world applicability and robustness. The results of this evaluation yielded a receiver operating characteristic (ROC) curve with an area under the curve (AUC) of 0.75. While not perfect, an AUC of 0.75 demonstrates that the model is capable of making reliable predictions, highlighting its potential as a valuable diagnostic tool.

Cross-dataset learning, in particular, offers great potential for enhancing the generalizability of miRNA biomarkers by enabling models trained on one population to be adapted for use in others. Although, to achieve broader generalizability, data from diverse populations and conditions will be needed. This would be a significant step toward standardizing miRNA biomarkers for clinical applications.

It has now been known that freely circulating blood miRNA may target other cells and organs to affect the expression of the cognate target gene. The results of our study provide significant insights into the biological functions and pathways regulated by the miRNAs we selected (miR-21-5p, miR-141-3p, and miR-221-3p). Through Gene Ontology (GO) the enrichment of biological processes (BP) is related to the regulation of nitrogen compound metabolism, macromolecules, and nucleobase compound metabolism underscores the involvement of these miRNA-targeted genes in fundamental cellular processes. Since cancer cells often exhibit altered metabolic pathways to sustain rapid proliferation, the association of our identified genes with metabolic regulation suggests that these miRNAs play a crucial role in the metabolic reprogramming characteristic of tumorigenesis. The connection to nucleobase compound metabolism, which is integral to DNA replication and repair, further highlights their potential contribution to genomic instability—a hallmark of cancer progression. KEGG pathway analysis provided further validation of the relevance of these miRNAs and their target genes in cancer. The enrichment in pathways such as miRNAs in cancer, TNF, JAK-STAT, MAPK, and Akt signaling highlights the involvement of our identified miRNAs in key regulatory networks that drive tumorigenesis, proliferation, and survival. The association with growth hormone and prolactin signaling pathways, which are known to be linked to PCa development and progression **(51)**, further strengthens the potential role of these miRNAs in the disease. The functional validation of the identified target genes remains open for future studies. Combining miRNA profiling with other omics data, such as proteomics or metabolomics, may further enhance the roles of these miRNAs.

Despite the promising results, our study has some limitations. The sample size, while sufficient for a preliminary study, needs to be expanded to validate our findings across a larger and more diverse cohort. Additionally, while whole blood provides a comprehensive miRNA profile, its complexity could introduce noise into the data, potentially obscuring cancer-specific miRNA signals. Finally, although ML improved the accuracy of our diagnostic model, further optimization is needed to ensure its reliability in clinical settings.

## 5. CONCLUSION

In conclusion, This study highlights the efficacy of integrating miRNA biomarkers with ML models to significantly enhance the diagnostic precision for PCa. By employing a Random Forest classifier, we were able to exploit complex interactions between miR-21-5p, miR-141-3p, and miR-221-3p, improving the model’s ability to differentiate PCa from BPH with increased sensitivity and specificity. The application of bioinformatic analyses, including functional enrichment and pathway interaction studies, further substantiated the critical roles of these miRNAs in PCa pathogenesis, underscoring their utility as reliable biomarkers. This approach established a ground framework for clinical diagnostics. Nonetheless, further validation across larger, multi-ethnic cohorts is required to enhance model generalizability and optimize performance for routine clinical implementation. This work provides a strong basis for advancing non-invasive, miRNA-based diagnostic solutions powered by ML in PCa management.

## Author contributions

*Research conceptualization and design*: Jain G; *Supervision and Validation*: Jain G; *Sample Collection and/or assembly of data*: Singh S and Kural S; *miRNAs analysis workflow:* Singh S; *Data analysis and interpretation*: Pathak AK, Singh S, Jain G, Gupta M and Kural S; *Visualization*: Singh S, Pathak AK and Kural S; *Writing-Original Draft*: Singh S, Pathak AK and Jain G; *Resources*: Yadav M and Bhardwaj MG; *Clinical Advisory:* Trivedi S, Kumar L and Yadav M; *Model optimization assistance:* Pathak AK and Gupta M *Writing-Review & Editing*: Singh S, Pathak AK, Kural S, Jain G, Trivedi S and Das P.

### Funding

This research was supported by the grant from BIRAC-India, under AGC-JanCare scheme to MIRNOW. The Institute of Eminence (IoE) Scheme at Banaras Hindu University, Fellowship/Grant support to Dr. Garima Jain, Banaras Hindu University. Non-Net fellowship support to AP and SK, and MIRNOW employee support to SS. We would also like to acknowledge the Centre for Genetic Disorders, BHU for providing laboratory resources and facilities as well as the Department of Urology, BHU for facilitating sample collection. We extend our sincere gratitude to all collaborators and patients involved in this study.

### Data Availability Statement

The data generated and analyzed in this study are available from the corresponding author upon reasonable request. To ensure privacy and comply with ethical guidelines, access to patient related data is restricted. However, anonymized datasets and supplementary materials used in this study will be provided on request for research and verification purposes, subject to institutional approval and appropriate data-sharing agreements.

