## Supplementary File for "miRNA Biomarkers in Prostate Cancer: Leveraging Machine Learning for Improved Diagnostic Accuracy"

### Supplementary File 1.

| miRNA name | Body Fluid | Expression Level in PCa | Function | Reference |
| --- | --- | --- | --- | --- |
| miR-21-5p | Plasma, Serum | Upregulated in PCa correlating with cancer progression | Promotes proliferation, migration, invasion, and chemoresistance via PTEN/AKT pathway. | (1–4) |
| miR-375-3p | Plasma, Serum | Increase in PCa compared to BPH | Enhances aggressiveness by regulating proliferation, migration, and metabolic pathways | (1,5,6) |
| miR-19B-3p | Serum, plasma | Elevated in metastatic PCa | Influences cell proliferation, migration, and metabolic pathways | (7,8) |
| miR-93-5p | Serum, Plasma | Upregulated in prostate cancer patients compared to healthy controls. | Promotes growth, invasion, and chemoresistance through TGF-β and PTEN/PI3K/AKTpathways | (8–10) |
| miR-141-3p | Serum, Plasma | Upregulated in prostate cancer patients compared to healthy controls and benign conditions. | Involved in epithelial-mesenchymal transition (EMT), metastasis. | (11–13) |
| miR-221-3p | Serum, Plasma | Elevated in Aggressive, advanced PCa | Supports tumor Growth,invasion, and metastasis via PI3K/AKT and cell cycle regulation | (14,15) |

***Supplementary Table. 1:*** *miRNAs promote prostate cancer progression, serving as potential biomarkers for tumor growth and metastasis.*

| Oligo Name | 5'<-----Sequence----->3' | Length |
|---|---|---|
| miR-21-5p(SL) | GTCGTATCCAGTGCAGGGTCCGAGGTATTCGCACTGGATACGACTCAACA | 50bp |
| miR-21-5p(FP) | ACCACCGTAGCTTATCAGACT | 21bp |
| miR-375-3P(SL) | GTCGTATCCAGTGCAGGGTCCGAGGTATTCGCACTGGATACGACTCACGC | 50bp |
| miR-375-3P(FP) | AACCGGTTTGTTCGTTCGG | 19bp |
| miR-19B-3P(SL) | GTCGTATCCAGTGCAGGGTCCGAGGTATTCGCACTGGATACGACTCAGTT | 50bp |
| miR-19B-3P(FP) | AACCGGTGTGCAAATCCATG | 20bp |
| miR-93-5P(SL) | GTCGTATCCAGTGCAGGGTCCGAGGTATTCGCACTGGATACGACCTACCT | 50bp |
| miR-93-5P(FP) | GGCCGCCAAAGTGCTGTT | 18bp |
| miR-141-3p(SL) | GTCGTATCCAGTGCAGGGTCCGAGGTATTCGCACTGGATACGACCCATCT | 50bp |
| miR-141-3P(FP) | AACCGGTAACACTGTCTGGT | 20bp |
| miR-221-3p(SL) | GTCGTATCCAGTGCAGGGTCCGAGGTATTCGCACTGGATACGACGAAACC | 50bp |
| miR-221-3p(FP) | AACACGCAGCTACATTGTCTG | 21bp |
| UniversalRevQ | CCAGTGCAGGGTCCGAGGTA | 50bp |
| RNU6 FP | CTCGCTTCGGCAGCACA | 21bp |
| RNU6 RP | AACGCTTCACGAATTTGCGT | 20bp |

***Supplementary Table 2:*** *Primer sequence details of all 6 miRNAs*

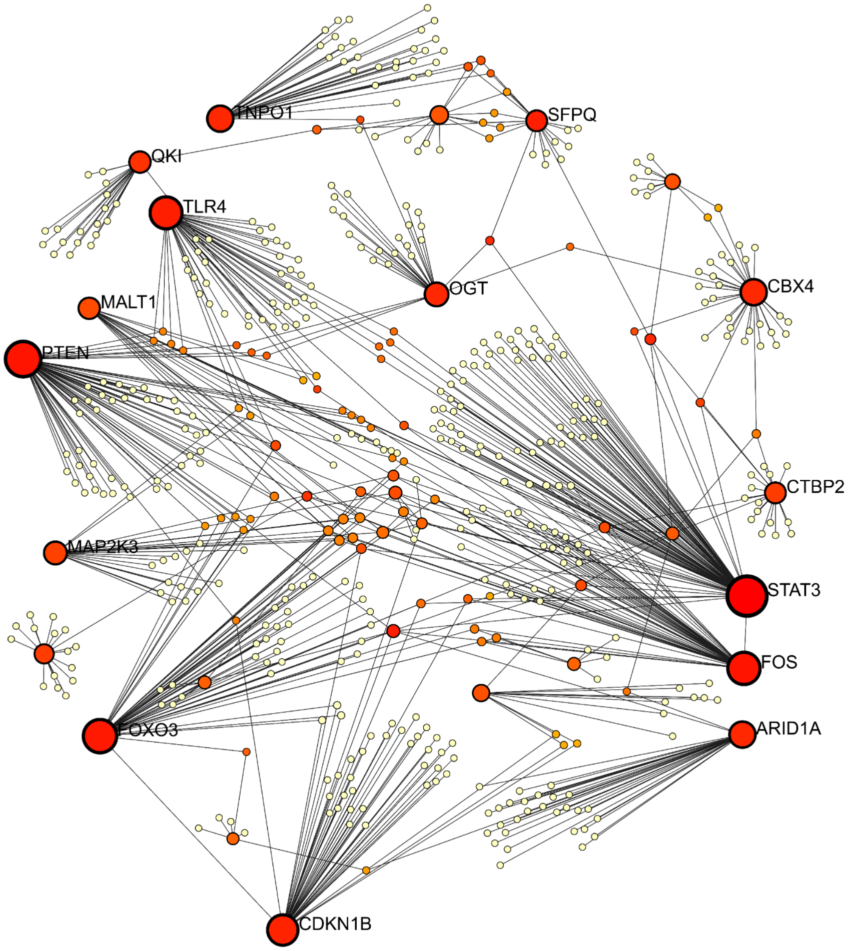


***Supplementary Fig. 1*** *: Protein-Protein Interaction Network of Key Genes Regulated by miRNAs in Prostate Cancer.*

***References***

12. Canatan D, Yılmaz Ö, Sönmez Y, Çim A, Baykara M, Savaş M, et al. Use of MicroRNAs as

biomarkers in the early diagnosis of prostate cancer. Acta Bio Medica Atenei Parm [Internet]. 2022 [cited 2024 Oct 14];93(3). Available from: https://www.ncbi.nlm.nih.gov/pmc/articles/PMC9335414/

13. Westermann AM, Schmidt D, Holdenrieder S, Moritz R, Semjonow A, Schmidt M, et al. Serum microRNAs as biomarkers in patients undergoing prostate biopsy: results from a prospective multi-center study. Anticancer Res. 2014;34(2):665–9.

14. Lyu J, Zhao L, Wang F, Ji J, Cao Z, Xu H, et al. Discovery and Validation of Serum MicroRNAs as Early Diagnostic Biomarkers for Prostate Cancer in Chinese Population. BioMed Res Int. 2019 Aug 25;2019:1–9.

15. Kawaguchi T, Komatsu S, Ichikawa D, Morimura R, Tsujiura M, Konishi H, et al. Clinical impact of circulating miR-221 in plasma of patients with pancreatic cancer. Br J Cancer. 2013;108(2):361–9.
